# CONCLAVE: CONsensus CLustering with Annotation-Validation Extrapolation for cyclic multiplexed immunofluorescence data

**DOI:** 10.1101/2025.11.13.688190

**Authors:** Pouya Nazari, Alexandre Arnould, Madhavi Dipak Andhari, Maite Fontecha, Jorge Martín Hernández, Bart De Moor, Jon Pey, Frederik De Smet, Francesca Maria Bosisio, Asier Antoranz

**Affiliations:** Translational Cell and Tissue Research Unit, Department of Imaging and Pathology, KU Leuven, Leuven, Belgium; The Leuven Institute for single-cell omics (LISCO), KU Leuven, Leuven, Belgium; The Leuven Cancer institute (LKI), KU Leuven, Leuven, Belgium; The laboratory for precision cancer medicine, Translational Cell and Tissue Research Unit, Department of Imaging and Pathology, KU Leuven, Leuven, Belgium; Center for Dynamical Systems, Signal Processing, and Data Analytics (STADIUS), Department of Electrical Engineering (ESAT), KU Leuven, Kasteelpark Arenberg, Leuven, Belgium; Intelligent Biodata Ltd, San Sebastian, Spain; Department of Pathology, UZ Leuven, Leuven, Belgium

## Abstract

High-dimensional cyclic multiplexed immunofluorescence (cMIF) enables single-cell phenotyping within intact tissues. Cell annotations rely on a multi-step pipeline involving normalization, sampling, dimensionality reduction, and clustering, but the absence of standardized benchmarks for method selection—especially at the clustering stage—leads to inconsistent and less reproducible phenotyping. To address this, we developed CONCLAVE, a consensus-clustering-based workflow that optimizes upstream steps and integrates results from multiple clustering algorithms retaining only those cell labels supported by at least two independent methods. Through in-silico simulations and real-world cMIF datasets, CONCLAVE consistently outperformed single-clustering-method approaches in accuracy, reproducibility, and robustness, with improvements becoming more evident when mapped within spatial tissue contexts. Additionally, CONCLAVE includes a scoring module that flags regions likely to contain unreliable or inconsistent data, facilitating targeted quality control. In summary, CONCLAVE offers a robust framework for cell annotation in cMIF datasets, enhancing the reliability of downstream spatial proteomics analyses.

**Graphical abstract:** 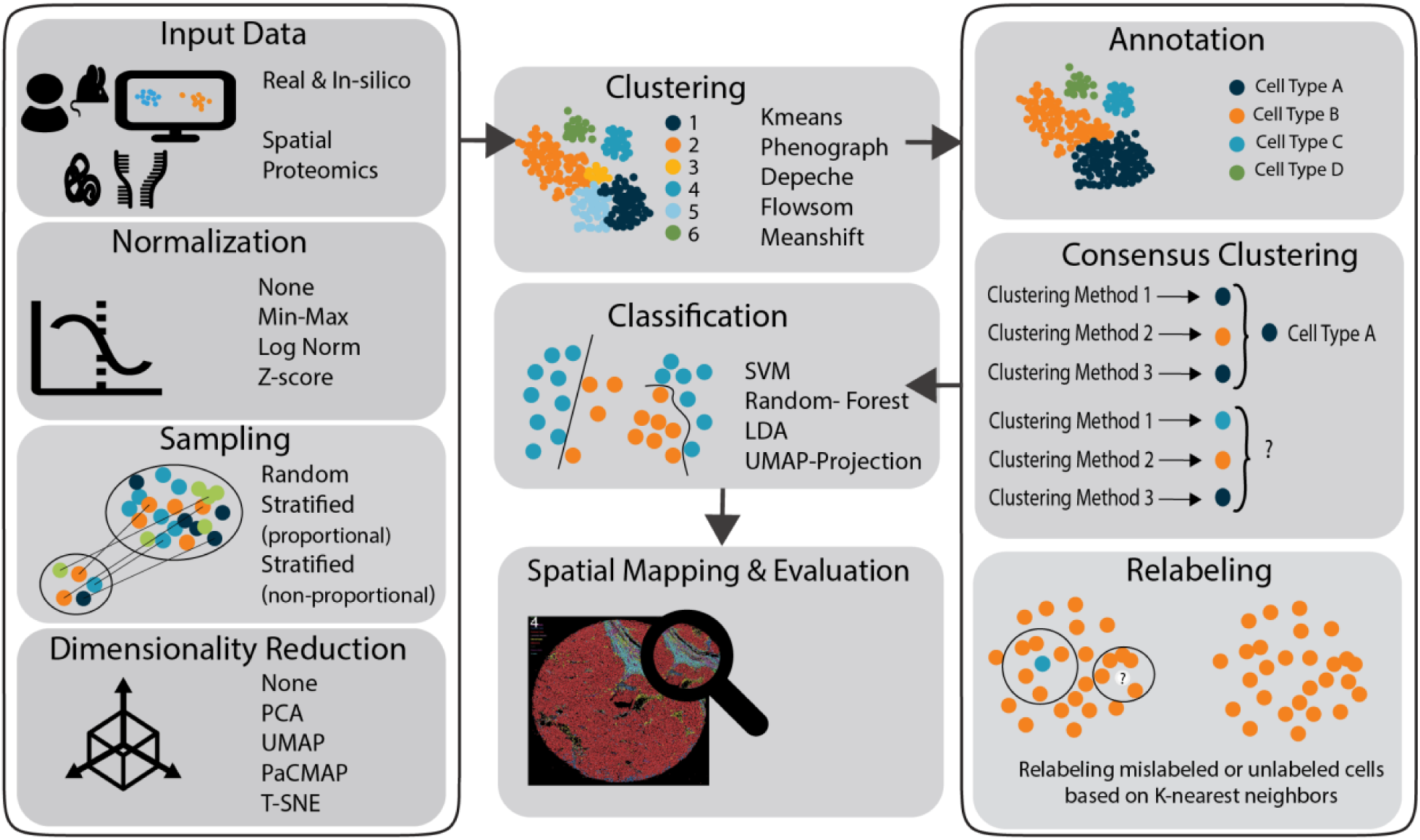

## Introduction

Traditional molecular profiling often sacrifices spatial information that is essential for understanding tissue organization and cell-cell interactions. Spatially resolved technologies, such as cyclic multiplexed immunofluorescence (cMIF), overcome this limitation by combining histological context with single-cell protein profiling. Measuring dozens of protein markers directly *in situ* allows cMIF to uncover phenotypic heterogeneity, microenvironmental structure, and cellular interactions that drive disease progression. [(Antoranz et al., 2022); (Vandereyken et al., 2023); (Lee et al., 2020)]. This preservation of spatial context is particularly valuable in oncology and immunopathology, where understanding the spatial arrangement of cells is essential for elucidating disease mechanisms [(Keren et al., 2018); (Schapiro et al., 2017)]. Furthermore, spatial data enables the identification of disease-specific structures, such as perivascular niches or tertiary lymphoid structures, which are not captured by dissociated single-cell approaches [(Jackson et al., 2020); (Moncada et al., 2020)].

Despite its obvious scientific advantages, the implementation of cMIF to evaluate real-work pathological samples comes with significant challenges, not in the least the computational analysis that is required to interpret the data. Unlike sequencing-based platforms, cMIF produces image-based data, introducing unique upstream sources of variability. Preprocessing steps such as image registration, and cell segmentation are essential for data quality and greatly impact the cellular quantifications and downstream steps [(Vandereyken et al., 2023); (Bérubé et al., 2022);(Weigert & Schmidt, 2022); (Stringer et al., 2021)]. Among these steps, cell phenotyping is critical to effectively profile and interpret tissue sections using cMIF, but this process is also affected by inherent biological confounding factors such as tissue heterogeneity and downstream computational aspects such as data normalization and integration. This becomes even more challenging when integrating data from different sources due to the lack of standardized computational workflows for cMIF (Chen et al., 2020), (Momenzadeh & Meyer, 2025). Historically, cell phenotyping was performed using manual gating, a strategy borrowed from flow cytometry, where cells were grouped into populations if the expression of given markers were higher/lower than a certain threshold. While effective for low-dimensional data, manual gating is labor-intensive, subjective, and poorly scalable for high-dimensional cMIF datasets [(Palit et al., 2019); (Lähnemann et al., 2020)] where thousands of images are often analyzed.

Computational approaches that are compatible with high-dimensional data can be broadly divided into supervised classifiers and unsupervised clustering methods. Supervised approaches, such as CellSighter (Amitay et al., 2023), rely on well-annotated datasets but struggle to generalize across tissues or detecting rare cell populations [(Bussi & Keren, 2024);(Lotfollahi et al., 2022)]. Unsupervised clustering methods, including Phenograph (Levine et al., 2015) and FlowSOM (Van Gassen et al., 2015), remain the gold standard as they provide flexibility and the potential to discover novel cell populations but are sensitive to preprocessing choices, parameter settings, and data quality, which can result in heterogeneous outputs [(Giesen et al., 2014); (Tan et al., 2020);(Kröger et al., 2024)].

Starting from cellular quantifications in tabular format following mean fluorescence intensity (MFI) extraction from cellular segmentation, a typical cell phenotyping pipeline includes several key steps: (1) normalization to correct technical variation introduced by antibody affinity (Heylen et al., 2024) or acquisition channel (Hahne et al., 2010), (2) sampling to manage the challenges of scalability and noise accumulation reducing computational load while preserving different cell types (Fan et al., 2014) (3) dimensionality reduction (DR) to retain critical features (Guldberg et al., 2023), (4) clustering to group similar cells (Ctortecka et al., 2024), and (5) cluster annotation based on canonical marker expression or expert knowledge (Lee et al., 2023). For each step, several alternatives are available, and it remains unclear which combination is optimal.

To address these challenges, we developed CONCLAVE, a consensus-driven framework designed for robust and reproducible cell phenotyping in spatial cMIF. By evaluating a wide range of normalization, dimensionality reduction, and clustering methods in both real and simulated datasets, we identified an optimal pipeline for cell classification. Importantly, by benchmarking CONCLAVE against current cell annotation pipelines on various data sets, we found that without consensus profiling, ∼20% of analyzed tissue samples contained regions with inconsistent or unreliable cell type annotations when mapped back into their spatial tissue contexts. By implementing a novel systematic relabeling approach, CONCLAVE markedly improved phenotypic coherence but also enabled us to implement a flagging system that highlights regions and/or samples with uncertain cell classifications, thereby streamlining targeted review and quality control. Together, these features establish CONCLAVE as a robust and accessible framework that enhances the reliability, interpretability, and transparency of spatial single-cell proteomics analyses, paving the way for more reproducible and biologically meaningful tissue profiling.

## Results

### Finetuning a multistep process using a ‘wisdom of the crowds’ approach across dummy and real-world cMIF data sets

Accurate cell annotation in spatial proteomics depends on a sequence of preprocessing and clustering steps, each of which can alter the outcome. As imaging data is highly susceptible to a myriad of artifacts (e.g., registration artifacts, segmentation artifacts, etc), cMIF is inherently noisy. As different preprocessing and clustering combinations can yield distinct, and sometimes conflicting, cell annotations, we hypothesized that individual algorithms make errors in different subsets of cells—each being partially correct. Consequently, integrating their outputs through a consensus, or “wisdom of the crowds,” strategy should offset method-specific biases and yield more robust and stable phenotyping results. To systematically evaluate this hypothesis, we combined more than 3,700 unique pipeline configurations, representing all possible combinations of normalization, sampling, dimensionality reduction, clustering, and classification strategies (see Table 1 for all included methods). This exhaustive analysis design enabled us to systematically benchmark how individual and combined analytical choices influence cell phenotyping accuracy. This was done using two in-silico (dummy_6 and dummy_25, see methods), and two publicly available real-world datasets. Next to analyzing the original datasets, we also introduced artificial perturbations (i.e. increasing the level of signal mixing in the in-silico datasets or by increasing the levels of stochastic noise in real datasets) to further map the features that mostly affect annotation performance (Fig. 1a-c). For all pipelines, we evaluated how individual analysis components influenced the recovery of predefined cell types (in-silico datasets) and annotated cell types (real datasets), but also how resilient each method combination was to noise.

**Figure 1.**
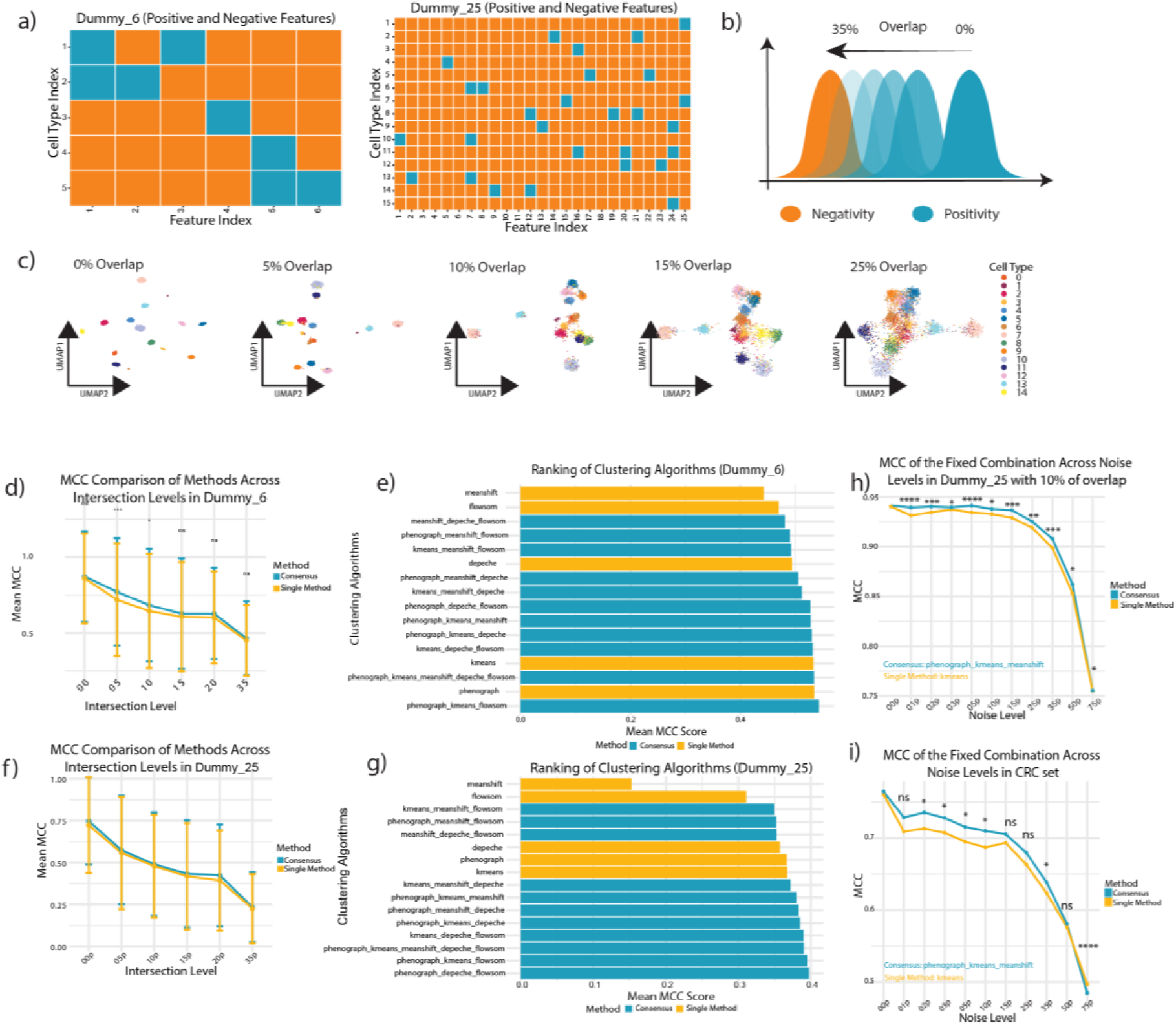
Design of dummy datasets and stability analysis a) Design of two synthetic benchmark datasets: dummy_6 with 6 features and 5 distinct cell-types and dummy_25 with 25 features and 15-cell types, with heatmaps of ground-truth feature profiles. b) Simulation of marker ambiguity via controlled feature overlap (0–35%) to mimic real-world mIHC challenges. c) Example clustering results for the dummy_25 dataset under 10% marker overlap, visualized with UMAP. d) MCC comparison for the 6-feature dataset; consensus clustering consistently outperforms individual methods across all overlaps. e) Ranking of method combinations for the 6-feature dataset; top-performing pipelines are dominated by consensus approaches (blue). f) MCC comparison for the dummy_25 dataset confirms consensus superiority in more complex datasets. g) Ranking of method combinations for the dummy_25 dataset; consensus pipelines remain highly stable across feature complexity. h) Robustness analysis under Gaussian noise (0–75%) on the 25-feature dataset; consensus clustering shows improved resilience compared to KMeans. i) Stability analysis in a real-world CRC dataset; consensus clustering maintains higher stability under increasing noise compared to KMeans alone.

**Table 1.** Advantages and disadvantages of the applied methods in different steps of the cell phenotyping pipeline. Summary of the key strengths and limitations of normalization, sampling, dimensionality reduction, clustering, consensus integration, and classification methods evaluated throughout this study.

| Step | Method | Pros | Cons |
| --- | --- | --- | --- |
| Normalization | None | Retains original data values. | Sensitive to variability in marker intensities across samples, which can lead to skewed results. |
|  | Min-Max | Scales data to a [0,1] range, easy to interpret and computationally efficient. | Can distort distribution if outliers are present. |
|  | Z-Score | Standardizes data, ensuring zero mean and unit variance. Useful for normally distributed data. | Sensitive to outliers, potentially skewing the normalized data. |
|  | Log-Normalization | Reduces the impact of large values, especially useful for highly skewed data distributions. | Not effective when data contains zeros, requiring additional transformations. |
| Sampling | Random Sampling | Simple and quick, avoids bias by selecting data points randomly. | May under-sample or over-sample important subpopulations, leading to an unbalanced representation. |
|  | Stratified Proportional Sampling | Ensures balanced representation by sampling proportionally from each tile in the tessellated UMAP space, preserving the original distribution of cellular phenotypes in a structured manner. | Can be computationally intensive; if the embedding or tiling does not perfectly reflect true biological diversity, underrepresented or noisy classes may still be under-sampled. |
|  | Stratified Non-Proportional Sampling | Enables targeted enrichment of rare or biologically relevant subpopulations by selectively oversampling from specific UMAP tiles. Useful when prior knowledge suggests importance of minority groups. | May introduce sampling bias if overrepresentation is not carefully controlled, potentially distorting population proportions and affecting downstream interpretation. |
| Dimensionality Reduction | PCA | Reduces dimensionality while preserving variance, fast and interpretable. | May not capture nonlinear relationships in complex spatial data. |
|  | t-SNE | Captures local structures and relationships, good for visualizing complex data clusters. | Computationally expensive and sensitive to parameter tuning, difficult to interpret globally. |
|  | UMAP | Preserves both local and global structures, faster than t-SNE, good for large datasets. | Sensitive to parameters; interpretability can be challenging for non-specialists. |
|  | PaCMAP | Preserves both global and local structures while being faster than t-SNE and UMAP. | Relatively new, limited standardization and fewer benchmarking studies for spatial data. |
| Clustering | Phenograph | Unsupervised, adaptable to diverse datasets, excels at detecting rare populations. | Slow with large datasets, sensitive to density-dependent clustering biases. |
|  | FlowSOM | Fast, robust, and effective in identifying rare populations, suitable for large datasets. | Requires expert knowledge for parameter tuning; results may vary based on initialization. |
|  | K-Means | Simple, interpretable, fast, effective when clusters are spherical. | Assumes equal-sized clusters; struggles with complex or imbalanced data distributions. |
|  | MeanShift | Does not assume a predefined number of clusters, can identify arbitrarily shaped clusters. | Computationally expensive, especially with large datasets, and sensitive to noise. |
|  | Depeche | Balances clustering with feature selection; reduces dimensionality while clustering, leading to interpretable results and automatic marker selection. | May oversimplify complex phenotypes; performance can degrade when relevant features are subtle or when rare populations are not well-separated. |

### Performance as a function of increasing mixing distributions

To systematically evaluate the robustness of our approach under varying data complexity and separability, we first applied it to two synthetic benchmark datasets, *dummy_6* and *dummy_25*. These datasets were designed to mimic single-cell expression profiles with controlled levels of marker overlap, allowing direct comparison of clustering performance as the signal-to-noise ratio changes. In the dummy_6 dataset, both consensus and single-method clustering exhibited a gradual performance decline as the overlap between cell populations increased, reflecting the expected loss of separability under higher signal mixing. Specifically, MCC values decreased from 0.872 ± 0.298 (consensus) and 0.860 ± 0.295 (single methods) at 0% overlap to 0.466 ± 0.243 and 0.453 ± 0.234 at 35% overlap, respectively. Although both approaches followed a similar decay trend, the consensus method consistently retained a modest advantage, on average +0.05 at 5% and +0.04 at 10% overlap (Fig. 1d,e). This difference was statistically significant at 5 % (adjusted p = 1.3 × 10⁻⁴), 10 % (adjusted p = 0.0496), and 20 % overlap (adjusted p = 0.0453; Wilcoxon test). In the dummy_25 dataset, the impact of increasing overlap was markedly stronger, with MCC values dropping by ∼69% across the 0–35% range for both approaches (from 0.748 ± 0.260 to 0.234 ± 0.208 for consensus; and from 0.723 ± 0.286 to 0.226 ± 0.206 for single methods, p = ns; Wilcoxon test). Importantly, the relative gap between consensus and single methods widened in this more complex data set, indicating improved resilience to noise of the consensus approach under higher dimensional data (Fig. 1f, g).

### Performance as a function of stochastic noise

We next assessed robustness under additive Gaussian noise (See Methods), a common source of variability in multiplexed imaging caused by weak or low staining quality. First, in dummy_25 (10% feature overlap), both single (KMeans) and consensus clustering (PhenoGraph + KMeans + FlowSOM) performed similarly on noise-free data (MCC = 0.940 vs 0.942). With increasing Gaussian noise, the consensus approach consistently retained higher accuracy. For example, at 25% noise, MCC was 0.900 for consensus vs 0.820 for KMeans (+0.08), and at 50% noise 0.863 vs 0.756 (+0.11) (Figure 1h). Differences were statistically significant across nearly all noise levels (adjusted p < 0.05 for 1 –50 % noise; Bonferrni-corrected Wilcoxon test), confirming that consensus clustering remains more robust as stochastic perturbation increases. We then applied the same analysis to a real-world cMIF dataset from colorectal cancer (CRC) (Lin et al., 2023). Here too, single and consensus methods were similar without additional noise (MCC = 0.761 vs 0.764, p = ns). As noise increased, consensus consistently outperformed KMeans, albeit with narrower margins: at 25% noise, 0.680 vs 0.663 (+0.017), and at 50% noise 0.581 vs 0.576 (+0.005) (Fig. 1i). These results confirm that consensus confers a measurable robustness advantage in noisy conditions, particularly in complex or synthetic settings. Significant differences were detected at moderate noise levels (2–10 % and 35 %, adjusted p < 0.05), and performance converged beyond 25 % (p > 0.05).

These results confirm that consensus clustering confers a measurable robustness advantage in noisy conditions, particularly in complex or synthetic settings. Importantly, as we describe in the sections below, the magnitude of the differences may be underestimated by other factors adding to the variability of the results (e.g., DR method, sampling approach).

### A systematic evaluation of the impact of preprocessing strategies on cell annotation accuracy

To establish best practices for cell phenotyping in noisy, high-dimensional spatial datasets, we systematically evaluated the performance of each preprocessing component (Table 1) using the *dummy_25* dataset (10 % feature overlap) and a real-world CRC dataset. Because rare cell types are often underrepresented and more susceptible to misclassification (Märtens et al., 2023), we examined two scenarios: (i) overall phenotyping accuracy across all cells and (ii) specific detection of rare populations, defined as those representing ≤ 5 % of the total cell population.

### Evaluation across all cell phenotypes

To identify optimal combinations, we applied recursive feature elimination (Fig. 2a; Supplementary Table 1). Beyond the exception of MeanShift, which consistently underperformed, clustering methods were not evaluated at this stage. This comprehensive screening revealed that not all methods for data preprocessing performed equally to reliable phenotyping. We first quantified the frequency of each method appearing within the top 10% and 25% of high-performing combinations (Fig. 2a). Methods such as LogNormalization, t-SNE, and MeanShift were rarely associated with high-scoring pipelines and were therefore excluded from subsequent analyses. Using the CRC dataset, we next compared method alternatives at each preprocessing step (Supplementary Fig. 1b–g), iteratively removing those that significantly underperformed. Evaluation of dimensionality-reduction (DR) methods showed that nonlinear techniques (UMAP and PaCMAP) introduced greater variance and substantial accuracy loss, particularly when combined with Linear Discriminant Analysis (LDA) for classification (Fig. 2b–c). Both methods performed significantly worse than no DR (Bonferroni-adjusted *p* = 8.9 × 10⁻³⁵ and 1.4 × 10⁻²³ for UMAP and PaCMAP, respectively) and PCA (*p* = 7.6 × 10⁻³¹ and 3.4 × 10⁻²¹, respectively; Wilcoxon rank-sum test). As these results exceeded the elimination threshold for this round (*p* ≤ 0.0001), UMAP and PaCMAP were removed from further analysis. In contrast, no DR performed marginally better than PCA (median MCC = 0.659 ± 0.113 vs. 0.649 ± 0.112; *p* = ns; Fig. 2b) and was retained for subsequent evaluations. At the normalization step, Z-score normalization yielded the highest median MCC (0.691 ± 0.084), outperforming MinMax (0.671 ± 0.125) and ‘no normalization’ (0.563 ± 0.098; *p* < 10⁻²⁰, Wilcoxon test; Fig. 2d). Finally, the choice of sampling strategy had a negligible impact, with no significant differences observed between random, stratified proportional, or stratified non-proportional approaches (Fig. 2e). Across all combinations, consensus clustering significantly outperformed single-method clustering approaches, achieving a higher median MCC (0.672 ± 0.068 vs. 0.624 ± 0.134; *p* = 1.3 × 10⁻¹², Wilcoxon test; Fig. 2f). Among clustering configurations, the PhenoGraph–K-Means–FlowSOM consensus achieved the best overall accuracy (0.687 ± 0.043), slightly surpassing the best single method, K-Means (0.682 ± 0.056; Fig. 2g). Classification performance was broadly comparable across methods, with Random Forest (median MCC = 0.664 ± 0.121), Support Vector Machine (SVM) (0.663 ± 0.121), and UMAP-based projection (0.661 ± 0.100) showing similar results, whereas LDA performed significantly worse (0.617 ± 0.102; Fig. 2h). Compared to LDA, Random Forest (*p* = 1.1 × 10⁻⁴), SVM (*p* = 6.9 × 10⁻⁵, and UMAP-projection (*p* = 1.5 × 10⁻⁵) all achieved significantly higher performance (Wilcoxon rank-sum test, Bonferroni correction).

**Figure 2.**
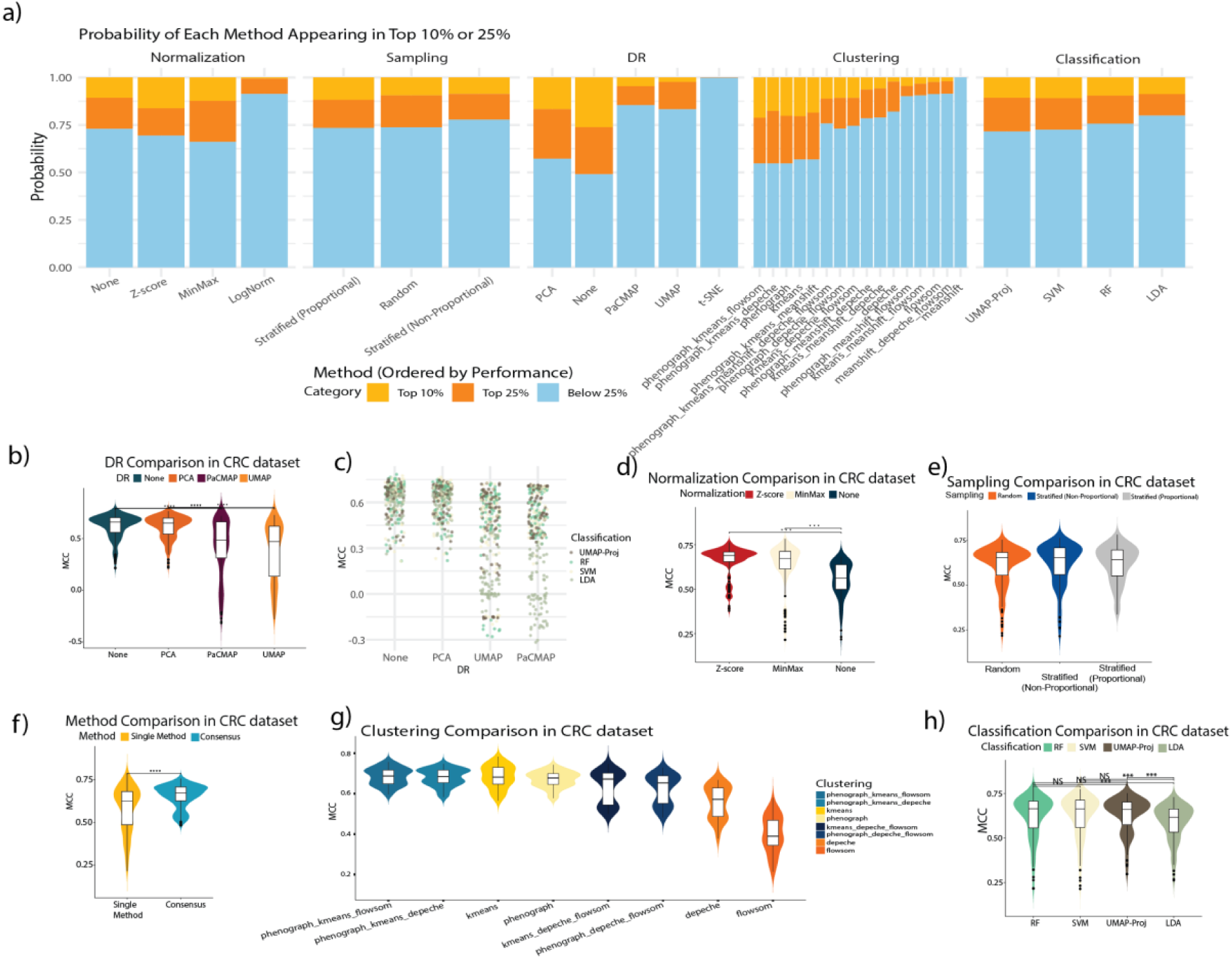
Evaluation of method combinations for cell phenotyping in spatial proteomics using dummy and CRC datasets for all cell types. **a)** Probability of each method appearing in the top 10%, top 25%, or below top 25% of performing pipelines. LogNormalization, t-SNE, and MeanShift appear most frequently in poorly performing combinations and are excluded from further evaluation. b) DR comparison reveals more stable performance for linear methods like PCA, while UMAP and PaCMAP occasionally underperform. c) MCC performance trends for different DR and classification combinations reveal that LDA is unstable when used with non-linear DR methods like PaCMAP or UMAP. d–h) Streamlined evaluation of selected methods on the CRC dataset (excluding t-SNE, PaCMAP, UMAP, LogNormalization, and MeanShift), illustrating the impact of each pipeline component: d) Z-score normalization remains the top performer. e) All sampling strategies yield similar performance, supporting inclusion in downstream evaluation. f) Consensus clustering again outperforms single methods. g) Clustering method comparison confirms that multi-method consensus outperforms individual clustering approaches. h) Final classifier comparison again shows LDA underperforms other tested approaches.

### Evaluation of rare cell phenotypes

As before, in the first round, underperforming methods (LogNormalization, t-SNE, MeanShift) were excluded. Across preprocessing steps, the results largely mirrored those observed for overall phenotyping accuracy (Supplementary Fig 2). In DR step, UMAP and PaCMAP once again underperformed no DR and PCA (Fig. 3b). Z-score normalization yielded the highest median MCC (0.384 ± 0.093), outperforming MinMax scaling (0.361 ± 0.095; Fig. 3c). Unlike the previous analysis, sampling strategy had a measurable impact on the identification of rare cell types (Fig. 3d). Stratified non-proportional sampling achieved the highest median MCC (0.398 ± 0.097), outperforming both random (0.369 ± 0.088; *p* = 8.6 × 10⁻⁴) and proportional stratified sampling (0.342 ± 0.094; *p* = 0.036; Wilcoxon test, Bonferroni correction). DR had no significant effect on rare-cell detection, with comparable median MCCs for no DR (0.659 ± 0.113) and PCA (0.649 ± 0.112; *p* > 0.05; Fig. 3e). Consensus and single-method clustering performed similarly in detecting rare populations (median MCC = 0.373 ± 0.057 and 0.375 ± 0.119, respectively; Fig. 3f), indicating no significant advantage of consensus integration for rare-cell detection. Among clustering strategies, K-Means achieved the highest performance for rare-cell detection (median MCC = 0.476 ± 0.059), outperforming the consensus combination of PhenoGraph–K-Means–FlowSOM (0.408 ± 0.044; Fig. 3g). This outcome was expected, as consensus integration in CONCLAVE prioritizes stable, reproducible labeling over maximizing sensitivity to rare subpopulations. Consequently, some rare cell types may not be fully preserved. This limitation can be mitigated through a hierarchical approach, where consensus clustering is first applied to define major cell types, followed by refined subclustering within each group using extended marker sets to recover rare subtypes more accurately. Finally, classification performance was comparable across Random Forest, SVM, and UMAP-projection, whereas LDA underperformed (Fig. 3h). Overall, considering both studies, the optimized CONCLAVE pipeline suggests Z-score normalization, stratified non-proportional sampling, no dimensionality reduction, and consensus clustering with PhenoGraph, K-Means, and FlowSOM, followed by UMAP-projection classification, as the best performing option. This configuration consistently outperformed alternatives across all analyses, offering superior normalization stability, improved recovery of rare cell types, and robust, interpretable classification with built-in confidence scoring (see Methods).

**Figure 3.**
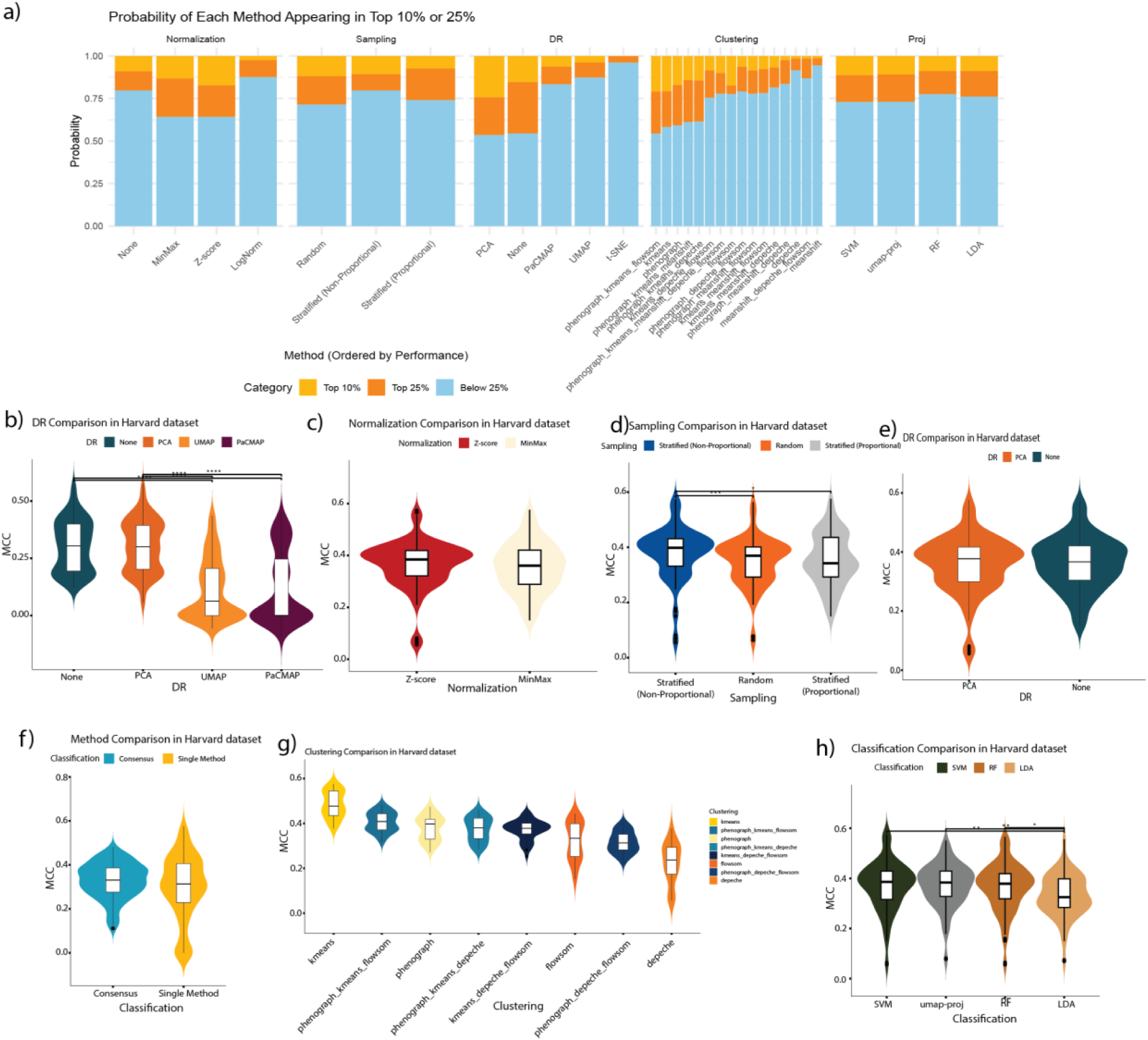
Evaluation of method performance for preserving rare cell types in the CRC dataset. **a)** Overall method performance across all pipeline steps. Bar plot showing the probability of each method appearing in the top 10% (orange) or top 25% (yellow) of combinations based on rare cell detection performance, across normalization, sampling, dimensionality reduction (DR), clustering, and classification steps. b) Impact of dimensionality reduction techniques. Avoiding dimensionality reduction entirely outperforms PCA, UMAP, and PaCMAP, indicating that DR may obscure critical local structures required for rare cell identification. c-h) Focused analysis after excluding underperforming methods (No normalization, UMAP, and PaCMAP): c) Z-score normalization outperforms MinMax in maintaining rare cell representation. d) Non-proportional stratified sampling yields better results for rare populations. e) Skipping dimensionality reduction provides the best outcomes. f) Consensus clustering shows more consistency in rare cell detection. g) Some single-method clustering approaches (e.g., KMeans) remain effective for subtypes. h) SVM and UMAP-based classifiers are more robust than LDA.

### Spatial Mapping Uncovers Misleading Cell-Type Assignments in Complex Tumors

After identifying the best-performing configuration, we evaluated CONCLAVE using an independent metastatic melanoma dataset (Antoranz et al., 2022). The analysis aimed to assess whether CONCLAVE maintains high phenotyping accuracy in new data, and to examine how different clustering approaches are capable of reconstructing tissue architecture in complex tumor microenvironments. Evaluating the outputs of individual clustering algorithms, only 54.9% of the cells were consistently classified with the same phenotype across all three methods, while 12.2% were differently labeled by the consensus approach (Fig. 4b). While individual clustering methods produced clusters with molecular profiles resembling known cell types, these were spatially inconsistent (Supplementary Fig. 3a). For instance, cells labeled as melanoma by FlowSOM were classified as dendritic by PhenoGraph and as epithelial by K-Means. These discrepancies indicate that different algorithms apply divergent decision boundaries even for well-separated phenotypes, leading to systematic mislabeling of distinct cell types. By integrating results through majority voting, CONCLAVE mitigates these method-specific biases, misclassifications that occur in different cell types across algorithms and producing more consistent annotations.

**Figure 4.**
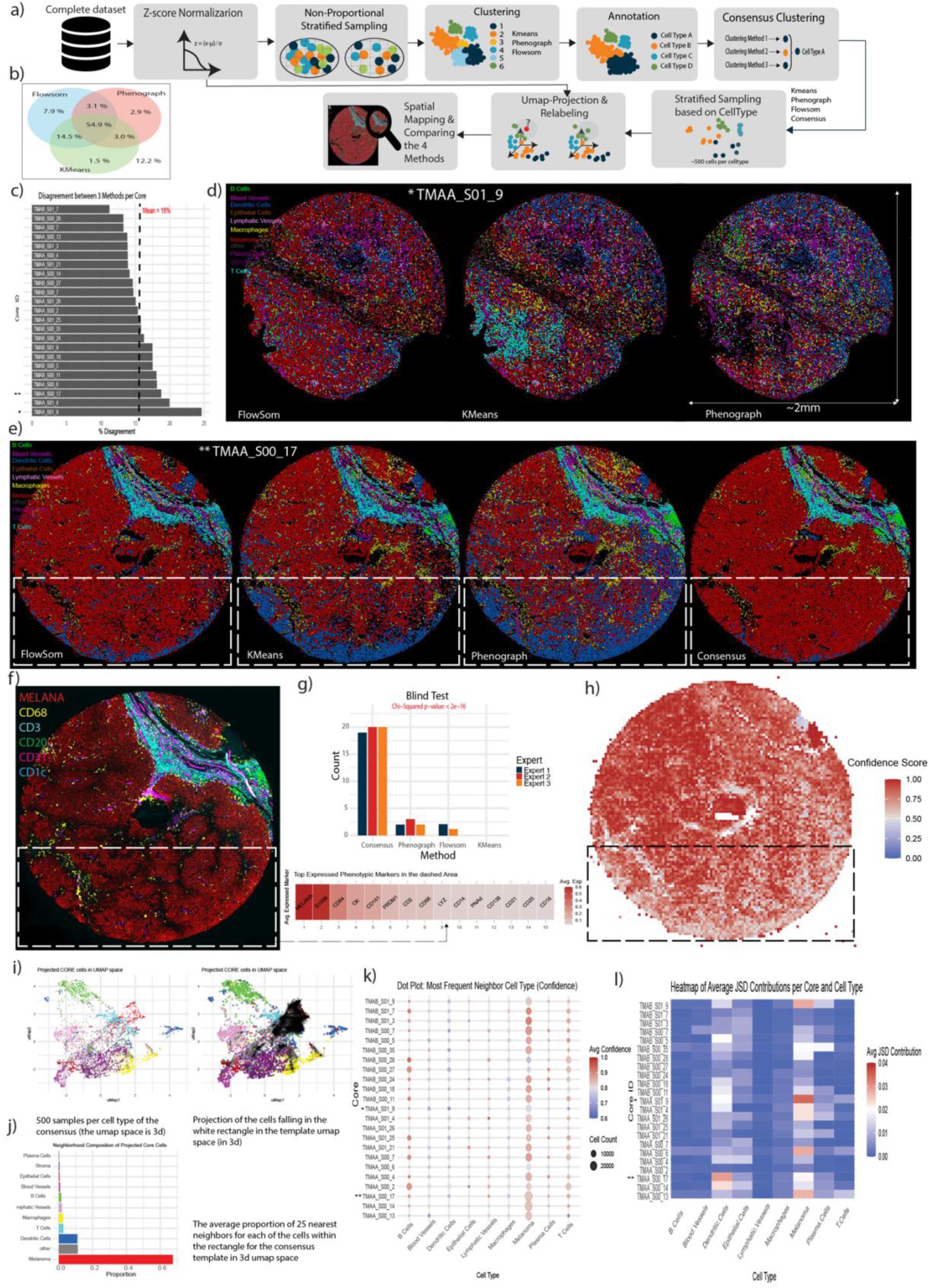
Application of the CONCLAVE pipeline to the Melanoma dataset and spatial validation of phenotyping robustness. **a)** Optimized pipeline for spatial proteomics phenotyping. Overview of the CONCLAVE workflow applied to the Melanoma dataset, incorporating z-score normalization, non-proportional stratified sampling, clustering with FlowSOM, KMeans, and PhenoGraph, followed by consensus integration, UMAP projection, and label propagation. b) Agreement between clustering methods and the consensus. Venn diagram showing the overlap between individual clustering methods and the consensus labels. A core 54.9% of cells are shared across all methods, while 12.2% are uniquely identified by the consensus. c) Disagreement score across tissue cores. Bar plot quantifying the percentage of disagreement between the three clustering methods per core. Cores with highest discordance are flagged for further analysis. d) Clustering result projections for a high-disagreement core. Spatial cell type maps from three individual clustering methods (FlowSOM, KMeans, PhenoGraph) in core TMAA_S01_9 show notable differences in regional labeling patterns. e) Comparison of clustering outputs and consensus. For core TMAA_S00_17, the spatial cell type distributions from each method and the consensus are shown. Within the dashed region, the consensus labels cells as Melanoma where individual methods classify them as Dendritic. f) Marker expression validation in ambiguous regions. Multiplex marker image (MelanA, CD68, CD3, CD20, CD21, CD1c) in the same core confirms the consensus label of Melanoma, supported by high MelanA and S100B expression in the boxed region. g) Expert blind evaluation of clustering accuracy. Bar plot summarizing expert pathologists’ preferences in a blind test across 23 cores. The consensus clustering result was most frequently chosen as the most biologically coherent (Chi-squared p < 2e–16). h) Single-cell confidence scores. Heatmap of confidence scores for core TMAA_S00_17. Lower scores in the boxed region correspond to cells with more uncertain identity, supporting use of this metric for quality control. i) UMAP projection of 500 cells per cell type. Left: Consensus labels in the shared UMAP space. Right: Projection of cells from the boxed region in (e), confirming the local enrichment of Melanoma cells in consensus-defined neighborhoods. j) Composition of neighborhoods in UMAP. Bar plot showing the dominant cell types in the nearest-neighbor neighborhoods of projected cells, with the consensus showing higher purity of Melanoma neighborhoods. k) Confidence scores per cell type across tissue cores. Dot plot showing the average confidence score for each cell type across all cores in the Melanoma dataset. Confidence is calculated based on the frequency of the most common neighboring cell type in the 3D UMAP space. Higher scores indicate greater neighborhood homogeneity and more stable phenotyping, particularly when cells are surrounded by neighbors of the same type. l) Heatmap of JSD contributions per core and cell type. Average Jensen–Shannon Divergence (JSD) contributions per cell type and core. Divergence is predominantly driven by Melanoma, Dendritic, and T cells, particularly in high-disagreement cores.

To assess which method best reflected biological reality, three expert pathologists independently performed a blind assessment of 23 tissue cores, comparing marker-stained images with clustering results from each method (Fig. 4f–g). The consensus approach received the majority of votes (59 of 69; *p* < 2 × 10⁻¹⁶, χ² test), indicating the closest alignment with histopathological interpretation. Among the three experts, FB conducted a more detailed evaluation that incorporated additional markers and spatial context, and her blind assessments were therefore used as the expert gold standard for quantitative comparison. Based on these annotations, the consensus method achieved the highest F1-score (0.903; Supplementary Fig. 3b), confirming its superior accuracy and biological coherence.

Mapping cell labels back to their spatial coordinates revealed that these discrepancies were not random but localized to specific tissue regions. Areas at tumor-immune interfaces or within dense infiltrates showed the greatest disagreement among methods. A disagreement score per core (see methods) identified regions of highest conflict (Fig. 4c–d). For example, in one representative core (Fig. 4e), individual methods predominantly labeled cells as dendritic cells, whereas the consensus correctly identified them as melanoma tumor cells.,These findings suggest that consensus clustering produces more reliable phenotypic templates for downstream classification, whereas individual clustering approaches generate less stable references that can propagate misclassifications during the projection of the full dataset.

### Quantifying Consensus: Spatial Coherence and Confidence Measures

To explore why the consensus method outperforms individual clustering algorithms, we assessed whether the improvement reflects the intrinsic stability of the consensus approach or is influenced by downstream relabeling. We implemented a spatial neighborhood validation to quantify local phenotypic coherence. In this analysis, we randomly sampled up to 500 cells per cell type and examined the 25 nearest neighbors of each cell in a three-dimensional UMAP embedding. Cells within the dashed region of the selected core (Fig. 4e) were projected into a shared UMAP space for each method (Phenograph, FlowSOM, KMeans, and consensus), and local consistency was quantified by calculating the proportion of neighboring cells that shared the same phenotype (Fig. 4i-j). This approach allowed us to directly compare the spatial coherence of cell type assignments across methods and determine whether the consensus clustering reduces local misclassification and noise inherent to individual algorithms.

This analysis revealed clear differences in local coherence between methods in dimensionality reduced space. In the examined core, cells identified as melanoma by the consensus were predominantly surrounded by other melanoma cells (Figs. 4i–j). In contrast, single-method results showed mixed neighborhoods with heterogeneous phenotypes (Supplementary Fig. 3c). These results highlight a key advantage of the consensus step: by excluding cells with conflicting cluster assignments, the consensus template is mathematically cleaner, containing fewer boundary or “edge” cells. This increases the separability of populations in feature space, yielding a more stable reference for the downstream projection and classification of all cells.

Despite the improved local coherence achieved by consensus clustering, some regions remained prone to uncertain classification. To quantify this, we introduced a single-cell phenotyping confidence score. For each cell, we calculated the proportion of its 25 nearest neighbors in the 3D UMAP space that shared the same label, producing a confidence value between 0 (no agreement) and 1 (complete agreement). Low-confidence scores indicate cells or regions where phenotyping is potentially unstable and more susceptible to misclassification (Fig. 4h). Applying this metric across all cores (Fig. 4k) allows systematic identification of regions with uncertain assignments, providing a practical tool for quality control. Importantly, this score also enables visualization of the effects of downstream relabeling, ensuring that spatial coherence improvements are quantifiable, and that subsequent analyses rely on the most reliable cell annotations. To further compare clustering outputs, we computed pairwise Jensen–Shannon Divergence (JSD) between cell-type distributions derived from the consensus and from each individual method across all tissue cores (Supplementary Fig. 3g; Fig. 4l). The resulting heatmap highlights which cell types contribute most to inter-method divergence. For example, in core TMAA_S01_9, differences were driven by melanoma, dendritic, and T-cell populations, whereas in core TMAA_S00_17, dendritic cells accounted for the largest deviation, consistent with prior spatial observations. High JSD values pinpointed tissue regions with substantial label disagreement, identifying phenotypes most affected by method-specific bias and highlighting samples that may benefit from targeted expert review.

Overall, applying the optimized CONCLAVE pipeline to the melanoma dataset demonstrates that consensus-based clustering enhances both the accuracy and spatial coherence of cell type annotation in complex tumor microenvironments. By integrating multiple clustering outputs, CONCLAVE mitigates method-specific biases and captures phenotypic patterns missed by individual algorithms, as evidenced by higher agreement with expert-curated labels, improved local neighborhood consistency, and superior F1-scores. The introduction of a cell-level confidence score and a flagging system further enables the identification of low-confidence or conflicting regions, providing a practical tool for quality control and ensuring reliable downstream analyses. Together, these results highlight that consensus clustering, complemented by targeted confidence measures, offers a robust and biologically interpretable framework for phenotyping cells in real-world spatial proteomics datasets.

## Discussion

Spatially resolved proteomics technologies such as cMIF provide invaluable insights into tissue cytometry and organization but require complex computational analyses. Accurate cell classification is essential for profiling these tissues and downstream spatial analyses, yet existing workflows suffer from variability introduced by different choices at the algorithmic level [(Vandereyken et al., 2023);(Bérubé et al., 2022);(Weigert & Schmidt, 2022)]. Manual gating can be applied in low dimensional settings, but it scales poorly with an increased number of markers. Supervised approaches can bridge this gap but are sensitive to overfitting and do not generalize robustly across cohorts. Unsupervised approaches such as clustering are sensitive to preprocessing choices and lack standardized benchmarks, leading to inconsistent or irreproducible results, particularly across rare populations (Chen et al., 2020). To address these challenges, we developed CONCLAVE, a consensus-based framework that integrates multiple clustering algorithms to achieve robust, reproducible, and spatially consistent cell phenotyping in multiplexed imaging datasets.

In CONCLAVE, we optimized upstream steps showing that the combination of Z-score normalization, non-proportional stratified sampling, and omission of dimensionality reduction prior to clustering yields the most accurate results in several in-silico and real cMIF datasets. At the clustering level, the consensus of PhenoGraph, K-Means, and FlowSOM outperformed any tested individual method, reducing instability particularly in heterogeneous or noisy tissue regions. Blind expert evaluation further confirmed that consensus-based phenotyping best captured histologically consistent cell populations. In addition, a quantitative quality-control metric was introduced to flag regions of uncertain classification, facilitating targeted review and human-in-the-loop validation. Together, these results demonstrate that CONCLAVE enhances the reliability and reproducibility of spatial proteomics phenotyping by stabilizing cluster definitions and minimizing algorithm-specific boundary effects.

Despite these advances, our study has several methodological and practical limitations. Methodologically, not all available options for each step were evaluated; future work should systematically test additional normalization, sampling, and clustering strategies as they continue to evolve. Practically, validation across a broader range of datasets and cMIF platforms (e.g., Comet, Phenocycler Fusion, Macsima) will be essential to assess the generalizability of our findings across acquisition technologies [(Vandereyken et al., 2023);(J.-R. Lin et al., 2018a);(Kinkhabwala et al., 2022)]. While CONCLAVE was compared against several unsupervised and supervised approaches, emerging methods based on latent mixed models (LMMs)(Hao et al., 2024), deep contrastive learning (e.g., CellCLIP, scJoint)(Y. Lin et al., 2022), or foundation models for multimodal single-cell analysis warrant future benchmarking. Finally, we have only tested CONCLAVE on feature matrices that capture the average expression of each cell (MFI) for cell phenotyping. Newer algorithms show increased cell profiling accuracy regardless of clustering approach by incorporating neighborhood context and spatial gradients into these feature matrices (e.g., UTAG, BANKSY, SpaGCN, GraphST) [(Kim et al., 2022);(Singhal et al., 2024);(Hu et al., 2021);(Long et al., 2023)]. CONCLAVE could synergize with these approaches towards more integrative representations and improved cell phenotyping. Additionally, while in this study CONCLAVE has been applied exclusively for cell phenotyping, the framework is equally compatible with any other task requiring clustering such as the classification of anatomical regions or the identification of cellular niches. The benefits added by consensus clustering in these scenarios remain to be explored. In brief, CONCLAVE establishes a consensus-based framework for *de novo* cell classification that optimizes preprocessing (normalization, sampling, and dimensionality reduction) and integrates multiple clustering algorithms through majority voting, providing a mathematically robust and reproducible resource for spatial proteomics analysis.

## Materials and methods

In this section, we present the CONCLAVE framework, the datasets used for benchmarking and validation, and the evaluation strategies applied to compare individual clustering methods against the consensus approach. The design of the Methods section follows the logical steps of the pipeline and mirrors the structure of the Results. First, we describe the datasets used to evaluate performance, including two controlled *in-silico* datasets and two real-world spatial proteomics datasets. Second, we outline the modular CONCLAVE pipeline, covering normalization, sampling, dimensionality reduction, clustering, annotations based on prior knowledge, consensus assesment, and relabeling/classification. Third, we detail the evaluation metrics and statistical analyses used to compare methods. Finally, we introduce the flagging system designed for quality control and uncertainty assessment.

### Datasets

To systematically evaluate phenotyping performance, we used two simulated (*in-silico*) datasets and two real-world spatial proteomics datasets. The *in-silico* datasets enabled benchmarking of clustering accuracy, stability, and robustness to noise under controlled conditions with known ground truth. The real-world datasets provided biologically relevant testbeds to assess generalizability across different tissues and marker panels.

### Simulated datasets

- **Dummy_6:** A simplified dataset of 50,000 cells described by 6 features and grouped into 5 cell types.
- **Dummy_25:** A more complex dataset of 50,000 cells with 25 features and 15 cell types. Each cell type expressed up to four features at high levels, allowing controlled overlap between classes.

For each dummy cell-type, we defined a positive/negative fingerprint of feature expression which was modelled using a positive and a negative Gaussian distribution with defined means and standard deviations. To simulate increasing classification difficulty, we varied the overlap between marker distributions from 0% to 35% (measured as shared area under the curves). Cell type proportions included both rare populations (<3% of total cells) and more dominant populations (5–40%). Figure 1a–c illustrates representative expression distributions and UMAP projections at different overlapping levels. These datasets provided the basis for benchmarking individual clustering methods and consensus strategies under controlled conditions.

### Real-world datasets

#### Colorectal Cancer (CRC)

We used the CRC002 whole-slide imaging sample from Lin et al. (2023), available through the public repository linked in the original publication. The dataset was acquired with the CyCIF technology (J.-R. Lin et al., 2018b) and comprised 483,936 cells, 21 cell types, and 24 markers. To focus on phenotyping, we restricted the marker panel to phenotypic and lineage-defining markers (Keratin, Ki67, PDL1, CD31, Desmin, aSMA, CD163, CD20, CD3, CD4, CD68, FOXP3, PD1, CD45, CD45RO, and CD8a). No further preprocessing was applied.

#### Melanoma

We used the TMA slide from Antoranz et al (2022), available through the public repository linked in the original publication. The dataset was acquired with the MILAN technology (Bolognesi et al., 2017) and comprised 542,637 cells, 19 cell types, and 36 phenotypic markers (*CD34, CD31, CD141, PNAd, CD25, CD14, CD1c, CK, CD21, FoxP3, CD23, GRB7, CD1A, Podoplanin, CD138, CD248, CD64, CD163, Pax5, IRF8, CD20, CD8, CD303, LYZ, CD16, CD2, HLADR, IRF4, CD5, CD79a, CD68, CD3, CD4, CD27, PRDM1, MELANA, S100B*). To simplify downstream analysis, several of the original cell types were consolidated into broader categories: Blood Vessels = Blood Vessels + High Endothelial Venules; Dendritic Cells = Follicular Dendritic Cells + Classical Dendritic Cells Type 1 (cDC1) + Classical Dendritic Cells Type 2 (cDC2) + Plasmacytoid Dendritic Cells (pDCs); Macrophages = M1-like Macrophages + M2-like Macrophages; T Cells = Regulatory T Cells + T Helpers + Cytotoxic T Cells. All other cell types (e.g., Melanoma, NK, B Cells, Plasma Cells, Langerhans Cells, Epithelial Cells, Lymphatic Vessels, Stroma) were retained as originally annotated. In both datasets, the input to CONCLAVE consisted of the per-cell feature matrices (cells as rows, markers as columns). Expression values represent marker intensities as defined in the original publications. No additional image-level corrections or marker scaling were applied, as analyses focused on the downstream phenotyping pipeline.

#### CONCLAVE Pipeline

In this study, we systematically benchmarked and optimized a modular spatial proteomics analysis pipeline, CONCLAVE, which integrates multiple preprocessing, clustering, and classification strategies. By adopting a consensus-based clustering approach, CONCLAVE combines the strengths of diverse methods to achieve more stable and accurate cell phenotyping, and its optimized configuration provides a generalizable framework for reproducible application to new datasets.

The CONCLAVE pipeline involves normalization, sampling, dimensionality reduction, clustering, annotation, relabeling and classification. Below we describe each step of the pipeline and the evaluated variants in detail.

#### 1. Normalization

We evaluated four normalization strategies applied on a per-sample basis to ensure comparability across samples while preserving within-sample marker distributions. For datasets consisting of a single sample (dummy datasets, CRC), normalization was applied once across the full dataset. For the melanoma dataset, which contained multiple samples, normalization was performed independently for each sample. The normalization strategies tested were:

- **Raw intensities:** features were used without transformation, preserving the original measurement scale.
- **Z-score normalization:** each feature was standardized to mean 0 and variance 1. To minimize the impact of extreme outliers, values were trimmed to the range defined by the 1st and 99th percentiles.
- **Quantile-based Min-Max scaling:** marker values were rescaled to the interval [0,1] based on the 1st and 99th percentiles, ensuring that rare outliers did not dominate the scaling.
- **Scaled logarithmic transformation:** a log transformation with a scaling factor was applied to reduce the dynamic range while preserving relative differences.

#### 2. Sampling

Sampling was introduced to reduce computational burden while ensuring that rare phenotypes and diverse regions of the feature space were represented (Fan et al. 2014). We implemented three strategies:

- **Random sampling:** cells were selected uniformly at random without stratification.
- **Stratified sampling (proportional):** cells were embedded into a 2D UMAP space derived from marker features, which was then divided into 16 strata (quartiles along each axis). Cells were sampled from each stratum in proportion to its size, preserving the natural distribution of the data.
- **Stratified sampling (non-proportional):** the same UMAP-based stratification was applied, but an equal number of cells was sampled from each stratum, ensuring balanced representation of the phenotypic landscape regardless of local density.

The reader should note that stratification in this step was performed on the UMAP feature space, not on annotated cell types. The goal at this step was to maintain coverage of the phenotypic landscape for clustering, not to artificially balance cell type frequencies.

#### 3. Dimensionality Reduction (DR)

**Dimensionality reduction (DR)** was applied to simplify the high-dimensional marker space while retaining biologically informative variation (Guldberg et al., 2023b). We tested five approaches:

- **No DR:** marker features were used directly.
- **Principal Component Analysis (PCA):** a linear projection capturing maximal variance (Bro & Smilde, 2014).
- **Uniform Manifold Approximation and Projection (UMAP):** a nonlinear method designed to preserve both local and global structure [McInnes et al., 2018].
- **Pairwise Controlled Manifold Approximation (PaCMAP):** a nonlinear method balancing local and global relationships [Huang et al., 2022].
- **t-distributed Stochastic Neighbor Embedding (t-SNE):** a nonlinear method primarily used for visualization [van der Maaten & Hinton, 2008], restricted to 3 dimensions.

To ensure fair comparison across methods, we restricted linear and nonlinear DR (except t-SNE) to 15 dimensions. This number was chosen to provide a consistent embedding size across methods while reflecting common practice in single-cell analysis workflows, where 10–30 dimensions are typically retained to capture major biological variation without overfitting to noise (e.g., default parameters in widely used tools such as Seurat).

#### 4. Clustering

Clustering was performed to generate candidate cell phenotypes using five commonly applied algorithms:

- **PhenoGraph**: a graph-based community detection method that identifies clusters as communities within a k-nearest neighbor graph [Levine et al., 2015].
- **KMeans**: a partitioning algorithm that assigns cells to k clusters by minimizing within-cluster variance [MacQueen, 1967]. To ensure comparability with other clustering methods, the number of clusters was set to the maximum number identified among the other algorithms (Phenograph and MeanShift) for each dataset.
- **FlowSOM**: a self-organizing map approach that captures both local and global data structures [Van Gassen et al., 2015]. We used the R implementation with default parameters, and the number of clusters was determined automatically by the method.
- **MeanShift**: a density-based algorithm that does not require specifying the number of clusters [Comaniciu & Meer, 2002].
- **DepecheR**: a probabilistic clustering framework robust to noisy data [Theorell et al., 2019].

#### 5. Annotation

Clusters are manually annotated using prior knowledge from domain experts to known cell phenotypes.

#### 6. Consensus Clustering

Consensus clustering was used to integrate results from multiple independent clustering algorithms into a single, robust cell type assignment.

- For each cell, the majority label across clustering methods was assigned.
- If no majority was reached (e.g. ties across methods), the cell was left temporarily unlabeled to avoid premature misclassification.
- In the optimized pipeline, consensus was performed across Phenograph, KMeans, and FlowSOM, which were selected after excluding underperforming methods.

This majority-vote strategy was chosen for its simplicity and interpretability, ensuring that cell types were only assigned when supported by at least two independent methods. By leaving ambiguous cells unlabeled at this stage, CONCLAVE preserved classification integrity and deferred uncertainty handling to the subsequent relabeling step (see below).

#### 7. Relabeling

Relabeling was performed to assign labels to ambiguous cells left unclassified after consensus clustering and to correct potential misclassifications. This step was applied only on the dataset obtained after Step 2 (sampling), not on the full dataset, to keep the process computationally efficient.

The procedure was as follows:

1. Class-balanced subsampling within the sampled dataset: from the already sampled dataset (Step 2), up to 500 cells per consensus-assigned class were selected to avoid class imbalance. This differs from the earlier feature-space stratified sampling in Step 2: here, subsampling was explicitly based on cell classes.
2. Dimensionality reduction: the balanced subset was projected into a 3D UMAP space to preserve local neighborhood structure.
3. K-nearest neighbor (KNN) relabeling: for each cell, the labels of its 25 nearest neighbors were considered.

a. Ambiguous (previously unlabeled) cells were assigned the majority class of their neighbors.
b. Confidently labeled cells were also included, which allowed correction of potential misclassifications.

This step ensured that all cells in the sampled dataset received consistent labels. In the final pipeline, these labels were then propagated to the full dataset during the classification stage.

## 8. Classification

Classification was the final step of the CONCLAVE pipeline, used to propagate labels from the sampled dataset to the full dataset. After relabeling ensured that all cells in the sampled dataset were consistently annotated, these labels served as the training set for supervised classifiers.

We evaluated several approaches:

- **Random Forest (RF):** an ensemble of decision trees that aggregates predictions to improve robustness [Breiman, 2001].
- **Support Vector Machine (SVM):** constructs optimal hyperplanes to separate classes in feature space [Cortes & Vapnik, 1995].
- **Linear Discriminant Analysis (LDA):** a linear method assuming normally distributed features with equal covariance [Fisher, 1936]. LDA consistently underperformed and was excluded from the final pipeline.
- **UMAP-based projection with KNN:** a simpler alternative in which cells from the full dataset are projected into a 3D UMAP embedding together with the sampled dataset, and labels are assigned based on the nearest neighbors. In this case, the relabeling and classification steps effectively merge into one procedure, since ambiguous cells are resolved and labels are propagated simultaneously.

The pipeline was first evaluated on the simulated datasets (dummy_6 and dummy_25) and the real CRC dataset to compare consensus versus single clustering methods under controlled conditions.

For benchmarking preprocessing and clustering strategies, we focused on the dummy_25 dataset with 10% feature overlap. This configuration was selected after visually comparing UMAP projections across overlap levels: lower overlaps yielded overly separated clusters that were unrealistically easy to resolve, whereas higher overlaps produced extensive blending that obscured population boundaries. The 10% overlap scenario provided an intermediate regime where clusters were still distinguishable yet sufficiently entangled to mimic the complexity observed in real mIHC datasets.

In the optimization stage, we applied the pipeline across over 3,700 unique methodological combinations, spanning different normalization, sampling, dimensionality reduction, clustering, consensus, and classification strategies. Both the dummy_25 (10% overlap) and the CRC dataset were used to identify consistently underperforming methods. The final optimization and method selection were performed using the CRC dataset alone, ensuring that the pipeline configuration was tailored to biologically complex, real-world data while still being informed by controlled in silico benchmarks.The resulting optimized workflow was then validated on the melanoma dataset to evaluate generalizability and robustness in an independent, heterogeneous tissue context.

### Evaluation Metrics

To assess clustering and classification performance, we used two complementary metrics:

- **Matthews Correlation Coefficient (MCC):** a measure of correlation between predicted and true labels, ranging from –1 (complete disagreement) to 1 (perfect prediction). MCC is well suited for multi-class classification and remains robust under class imbalance [Matthews, 1975; Chicco & Jurman, 2020]. We used the multiclass generalization of MCC as implemented in scikit-learn, which computes correlation across all classes simultaneously [Gorodkin, 2004; Chicco & Jurman, 2020]. For a confusion matrix C of size *K × K*, where *K* is the number of classes, the multiclass MCC is defined as:

where:

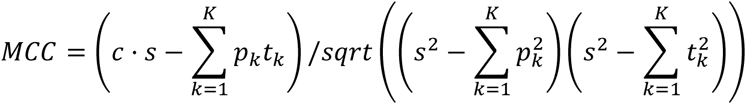

*C* = confusion matrix of size *K* × *K*

*c* = ∑_*i*_ *C*_*ii*_ (sum of diagonal = number of correctly predicted samples)

*p*_*k*_ = ∑_*i*_ *C*_*ik*_ (predicted for class *k*)

*t*_*k*_ = ∑_*i*_ *C*_*ki*_ (true samples of class *k*)

*s* = ∑_*i*,*k*_ *C*_*ik*_ (total number of samples)

**F1-score:** the harmonic mean of precision and recall. We report the macro-averaged F1-score across classes, ensuring that both abundant and rare cell types contribute equally to the metric.

### Statistical Analysis of Preprocessing Steps

To evaluate the effect of preprocessing steps (normalization, sampling, dimensionality reduction, clustering, classification), we compared MCC values across methods using pairwise Wilcoxon rank-sum tests. P-values were adjusted for multiple comparisons using the Bonferroni procedure. In visualizations (e.g., violin plots), statistical significance was indicated as follows: *** (p ≤ 0.0001), *** (p ≤ 0.001), ** (p ≤ 0.01), * (p ≤ 0.05), and NS (not significant). Significance annotations were omitted for clustering comparisons to prevent overcrowding of figures.

### Robustness Evaluation

To assess the stability of our phenotyping pipeline under noisy conditions, we introduced controlled Gaussian noise into both the dummy_25 dataset (10% overlap) and the CRC dataset.

For each marker, random noise was drawn from a normal distribution with mean zero and a standard deviation proportional to the marker’s observed variability. This ensured that the magnitude of perturbation scaled with the natural dynamic range of each marker, reflecting realistic sources of measurement noise in imaging-based datasets (*Chevrier et al., 2018); (Vu et al., 2022)*.

We applied ten increasing noise levels (1%, 2%, 3%, 5%, 10%, 15%, 25%, 35%, 50%, and 75% of the marker’s standard deviation). Each condition was repeated 30 times with different random seeds to account for stochastic variation.

Both noisy and original datasets were processed through the same pipeline. We compared KMeans as a representative single-method clustering and the consensus of Phenograph, KMeans, and Meanshift. These two setups were chosen as representative comparators because, in the absence of added noise, their MCC performance was closely matched (Figure 1h&i). This ensured that differences observed under increasing noise levels reflected robustness to perturbation rather than baseline discrepancies.

### Optimization of the pipeline

After benchmarking individual steps, we sought to identify the most robust overall pipeline configuration. To this end, we systematically evaluated ∼3,700 methodological combinations, spanning different choices of normalization, sampling, dimensionality reduction, clustering, consensus, and classification.

- **Initial screening:** both the dummy_25 dataset (10% overlap) and the CRC dataset were used to rank methods by performance. This stage allowed us to identify consistently underperforming approaches across controlled and real-world conditions.
- **Method exclusion:** based on this screening, LogNormalization, t-SNE, and MeanShift were excluded from further analysis, as they consistently ranked among the lowest-performing configurations (Figure 2b and Figure 3b).
- **Final optimization:** after excluding these methods, the CRC dataset was used exclusively to refine the pipeline. We compared remaining strategies across all modules using recursive elimination, progressively narrowing down to the most stable combinations. To ensure robustness, optimization was performed both considering all cell types and focusing specifically on rare populations (see results).

The optimized pipeline was subsequently applied to the independent melanoma dataset to evaluate generalizability. In this step, the mapped results were also inspected closely to confirm that the optimized workflow produced biologically coherent and spatially consistent annotations.

### Flagging System

In addition to consensus clustering, CONCLAVE includes a flagging system designed to highlight regions or samples with uncertain or inconsistent classifications. This quality-control layer enables users to identify tissue areas where clustering results may be unreliable and warrant closer inspection. Three complementary approaches were implemented:

#### 1. Disagreement Score

To quantify method disagreement at the core level, we defined a per-cell absolute no-consensus indicator and summarized it within each core. For each cell *i*i, let *Li={ℓi(1),ℓi(2),ℓi(3)}* be the labels assigned by PhenoGraph, FlowSOM, and KMeans. We set

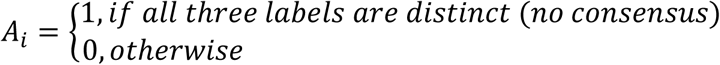

Cells with any blank label were not counted as absolute no-consensus.

For each core *c* with *Nc* cells, the disagreement score was computed as the percentage of absolute no-consensus cells:

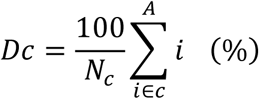

This metric highlights cores where clustering methods diverge strongly, while *Dc* indicates a larger fraction of cells where the three methods assign mutually different labels (Fig. 4c–d).

#### 2. Confidence Scoring

During the relabeling step, we estimated classification certainty for each cell based on neighborhood agreement in UMAP space. For each cell, its 25 nearest neighbors were identified, and the confidence score was defined as the proportion belonging to the majority class:

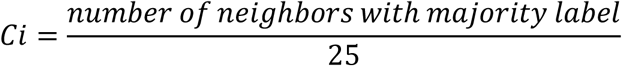

Confidence scores range from 0 (completely uncertain) to 1 (highly confident). These scores were visualized as spatial heatmaps (50 × 50-pixel tiles), with warmer colors indicating higher certainty (Fig. 4h). Average confidence per cell type was summarized in dot plots for different cores (Fig. 4k).

#### 3. Jensen–Shannon Divergence (JSD)

To assess differences between the consensus and individual clustering methods, we computed the Jensen–Shannon divergence (JSD), a symmetric measure of dissimilarity between probability distributions. For each cell:

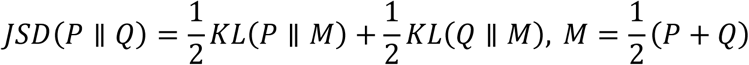

where *P* is the consensus label distribution and *Q* is the distribution from a single method. At the cell-type level, we calculated:

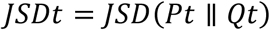

for subsets of cells of type *t*, and then computed each type’s contribution to overall divergence:

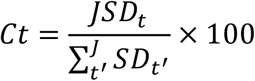

This highlighted which cell types were most responsible for differences between methods (Fig. 4l). At the core level, average JSD across method pairs was summarized, flagging cores with high disagreement (Supplementary Fig. 3g).

Together, these three flagging strategies — disagreement scores, confidence heatmaps, and JSD — provide complementary perspectives on classification reliability. They enable users to detect both global inconsistencies (across methods) and local uncertainties (within tissue regions), making CONCLAVE not only a consensus-based phenotyping pipeline but also a transparent quality-control framework.

---

**Supplementary Figure 1.**
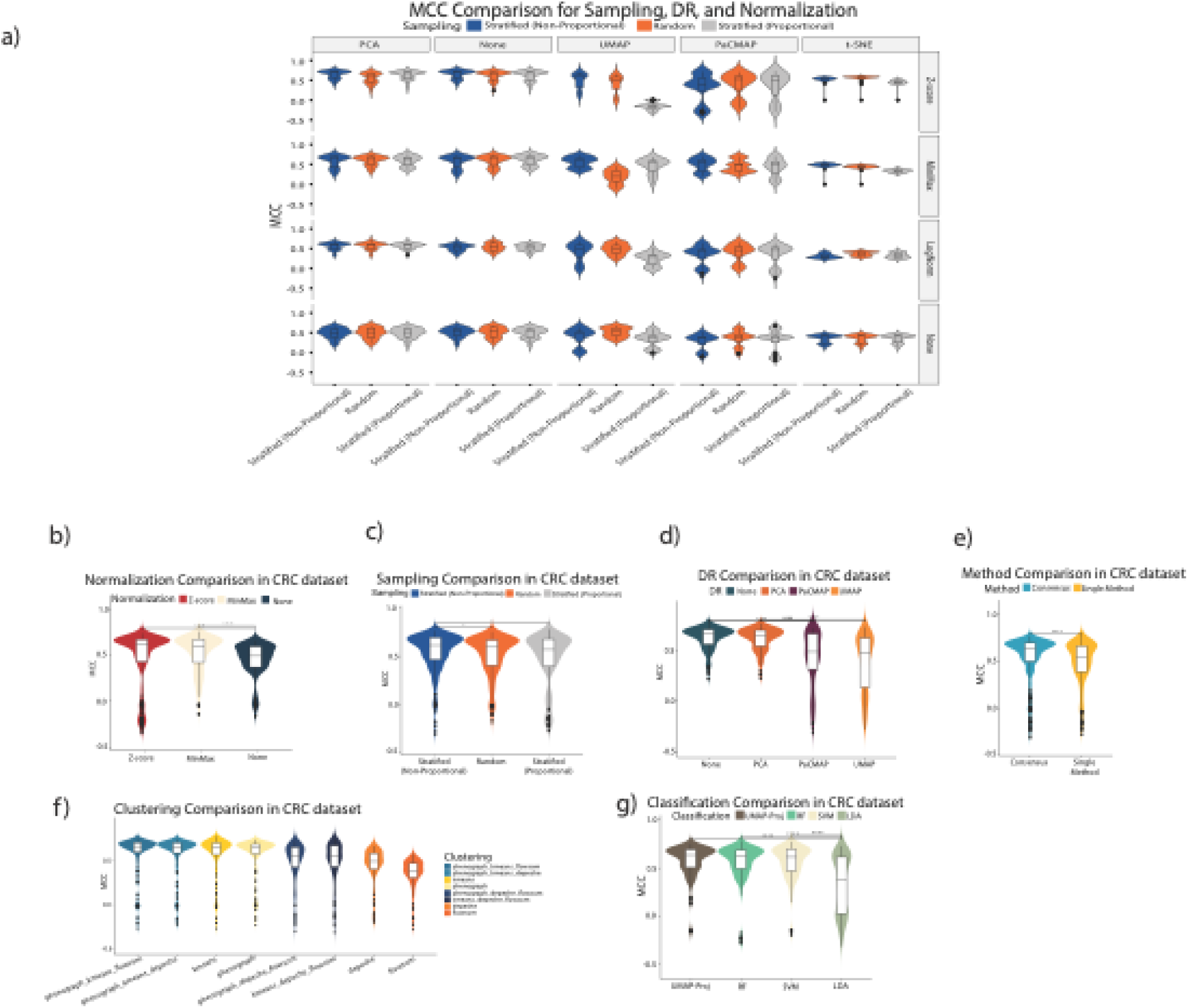
Evaluation of method combinations for cell phenotyping in spatial proteomics using CRC datasets considering all cell types. a) MCC performance comparison across all combinations of sampling strategies, normalization techniques, and dimensionality reduction (DR) methods using the CRC dataset. t-SNE consistently underperforms, and UMAP shows high variability. b-g) Evaluation of remaining methods on the CRC dataset after removing underperforming methods in round 1: b) Comparison of normalization strategies shows that Z-score normalization consistently outperforms MinMax and no normalization. c) Sampling strategy comparison shows minimal impact, with slightly better performance from stratified (non-proportional) sampling. d) DR comparison reveals more stable performance for linear methods like PCA, while UMAP and PaCMAP occasionally underperform. e) Consensus clustering significantly outperforms single-method clustering. f) Comparison of clustering methods shows improved stability and performance for consensus methods like phenograph_kmeans_flowsom and phenograph_kmeans_depeche. Single methods such as FlowSOM and depeche consistently underperform. g) Classification method comparison highlights LDA with more variability.

**Supplementary Figure 2.**
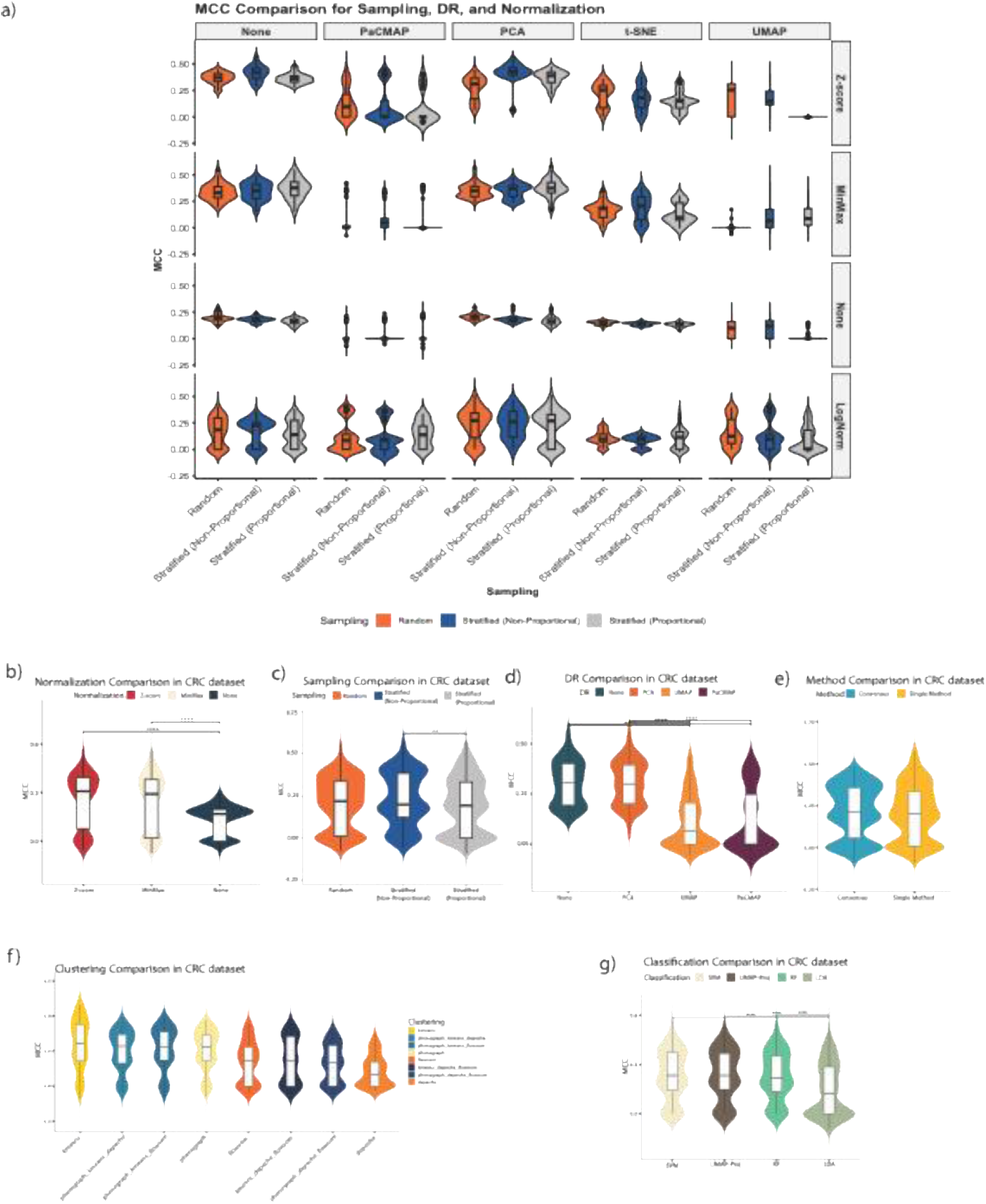
Evaluation of method performance for preserving rare cell types in the CRC dataset. a) MCC performance comparison across all combinations of sampling strategies, normalization techniques, and dimensionality reduction (DR) methods using the CRC dataset. t-SNE consistently underperforms, and UMAP shows high variability. b-g) Evaluation of remaining methods on the CRC dataset after removing underperforming methods in round 1 for rare cell types: b) Z-score normalization significantly outperforms MinMax scaling and the absence of normalization, highlighting its importance in preserving rare populations. c) Comparison of sampling strategies. Stratified non-proportional sampling shows improved performance in preserving rare cell types compared to random or proportional stratified sampling. d) Impact of dimensionality reduction techniques. Avoiding dimensionality reduction entirely outperforms PCA, UMAP, and PaCMAP, indicating that DR may obscure critical local structures required for rare cell identification. e) Consensus methods show superior overall performance over individual methods. f) Performance of individual clustering algorithms. Although consensus-based clustering generally performs better, some individual algorithms like KMeans perform competitively, particularly for rare subtypes. g) Classification algorithm performance. Support Vector Machines (SVM) and UMAP projection-based classifiers demonstrate better rare cell preservation compared to Random Forest (RF) and LDA.

**Supplementary Figure 3.**
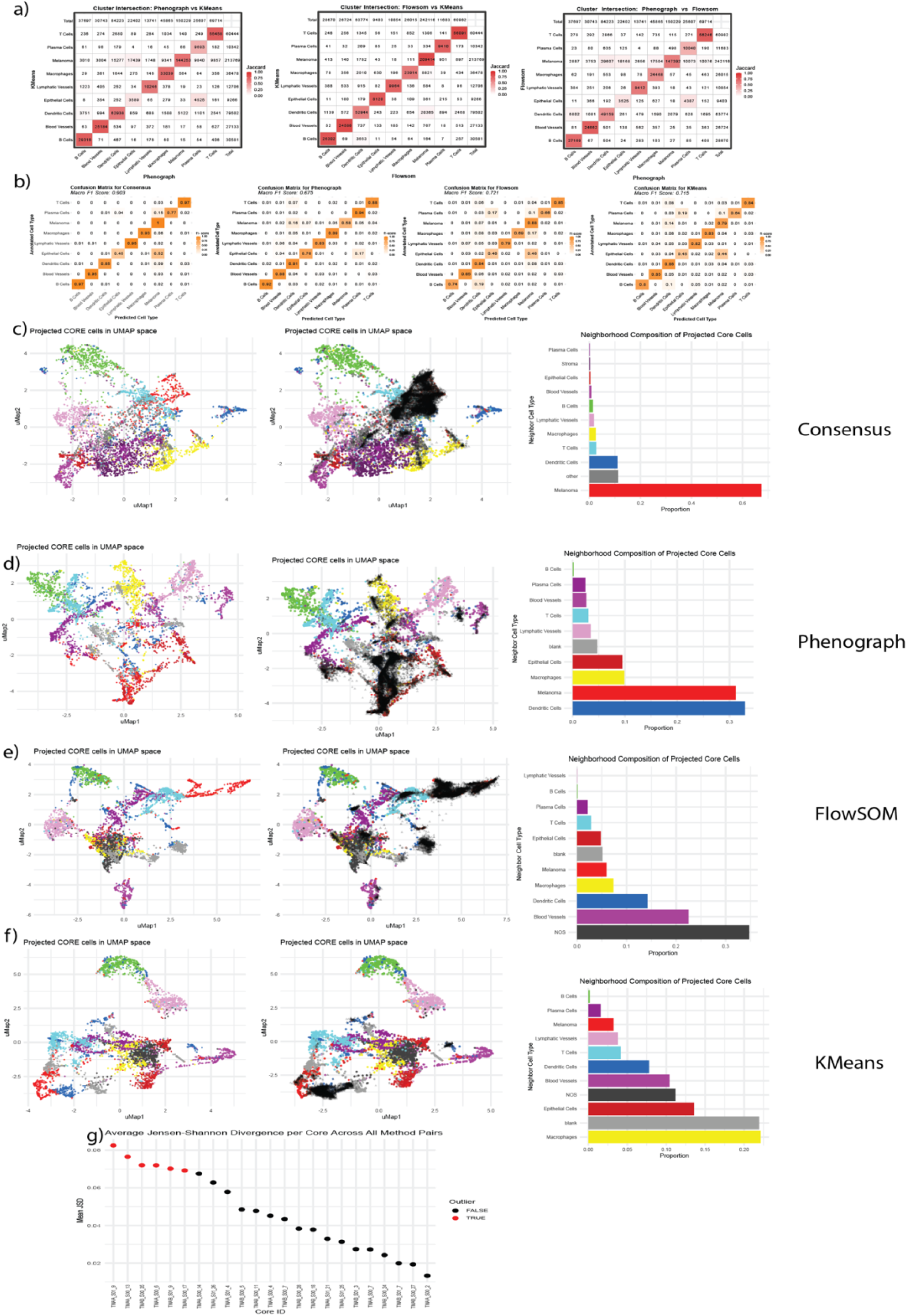
Comparative assessment of clustering methods in the Melanoma dataset using projection, spatial consistency, and divergence metrics. a) Pairwise cluster overlaps between individual methods. Heatmaps showing cluster-wise intersection between Phenograph vs. KMeans, FlowSOM vs. KMeans, and Phenograph vs. FlowSOM. Values represent the number of shared cells, highlighting inconsistent cluster mappings across methods. b) F1-score evaluation against expert-curated ground truth. Bar plot showing F1-scores for each clustering method using ground truth labels curated by the expert who thoroughly cross-validated marker expression across all tissue cores. The consensus method achieves the highest F1-score (0.903), significantly outperforming individual methods and demonstrating superior accuracy in resolving complex cell-type identities. c–f) Neighborhood consistency of projected core cells in UMAP space. Left and center: UMAP projections of 500 cells per cell type (colored background) with black points representing cells from the high-disagreement region in core TMAA_S00_17. Right: Bar plots showing the composition of the 25 nearest neighbors for these projected cells, stratified by cell type, for the Consensus (c), Phenograph (d), FlowSOM (e), and KMeans (f) outputs. The consensus maintains higher neighborhood purity of Melanoma cells, while single-method results show greater heterogeneity. e) Jensen–Shannon Divergence (JSD) across tissue cores. Bar plot summarizing average JSD scores per core across all pairwise method comparisons. Red dots indicate outlier cores with substantially higher divergence, often associated with noisy or ambiguous tissue regions. These flagged cores may warrant closer inspection or reannotation.

**Supplementary Table 1.** Feature elimination/selection in multiple rounds in CRC dataset considering all the cell types.

| Round | Removed/Selected Methods |
| --- | --- |
| 1 | Removed: LogNorm, t-SNE, Meanshift |
| 2 | Removed:UMAP, PaCMAP |
| 3 | Selected:<br>Z-score<br>Any sampling method<br>No DR<br>KMeans<br>Consensus of KMeans, Phenograph and FlowSOM<br>umap_projection |

**Supplementary Table 2.** Feature elimination/selection in multiple rounds in CRC dataset considering rare cell types.

| Round | Removed/Selected Methods |
| --- | --- |
| 1 | Removed: LogNorm, t-SNE, Meanshift |
| 2 | Removed: UMAP, PaCMAP |
| 3 | Selected:<br>Z-score<br>Stratified non-proportional sampling<br>No DR<br>KMeans<br>Consensus of KMeans, Phenograph<br>and FlowSOM<br>umap_projection |

## References

1. Bérubé, S., Kobayashi, T., Wesolowski, A., Norris, D. E., Ruczinski, I., Moss, W. J., & Louis, T. A. (2022). A pre-processing pipeline to quantify, visualize, and reduce technical variation in protein microarray studies. Proteomics, 22(3), e2100033. 10.1002/pmic.202100033

2. Bolognesi, M. M., Manzoni, M., Scalia, C. R., Zannella, S., Bosisio, F. M., Faretta, M., & Cattoretti, G. (2017). Multiplex Staining by Sequential Immunostaining and Antibody Removal on Routine Tissue Sections. The Journal of Histochemistry and Cytochemistry: Official Journal of the Histochemistry Society, 65(8), 431–444. 10.1369/0022155417719419

3. Chen, C., Hou, J., Tanner, J. J., & Cheng, J. (2020). Bioinformatics Methods for Mass Spectrometry-Based Proteomics Data Analysis. International Journal of Molecular Sciences, 21(8), 2873. 10.3390/ijms21082873

4. Ctortecka, C., Clark, N. M., Boyle, B. W., Seth, A., Mani, D. R., Udeshi, N. D., & Carr, S. A. (2024). Automated single-cell proteomics providing sufficient proteome depth to study complex biology beyond cell type classifications. Nature Communications, 15(1), 5707. 10.1038/s41467-024-49651-w

5. Fan, J., Han, F., & Liu, H. (2014). Challenges of Big Data analysis. National Science Review, 1(2), 293–314. 10.1093/nsr/nwt032

6. Guldberg, S. M., Okholm, T. L. H., McCarthy, E. E., & Spitzer, M. H. (2023). Computational Methods for Single-Cell Proteomics. Annual Review of Biomedical Data Science, 6(1), 47–71. 10.1146/annurev-biodatasci-020422-050255

7. Hahne, F., Khodabakhshi, A. H., Bashashati, A., Wong, C., Gascoyne, R. D., Weng, A. P., Seyfert-Margolis, V., Bourcier, K., Asare, A., Lumley, T., Gentleman, R., & Brinkman, R. R. (2010). Per-channel basis normalization methods for flow cytometry data. Cytometry Part A, 77A(2), 121–131. 10.1002/cyto.a.20823

8. Hao, M., Gong, J., Zeng, X., Liu, C., Guo, Y., Cheng, X., Wang, T., Ma, J., Zhang, X., & Song, L. (2024). Large-scale foundation model on single-cell transcriptomics. Nature Methods, 21(8), 1481–1491. 10.1038/s41592-024-02305-7

9. Heylen, D., Pusparum, M., Kuliesius, J., Wilson, J., Park, Y.-C., Jamiołkowski, J., D’Onofrio, V., Valkenborg, D., Aerts, J., Ertaylan, G., & Hooyberghs, J. (2024). Synthetic plasma pool cohort correction for affinity-based proteomics datasets allows multiple study comparison. Briefings in Bioinformatics, 26(1). 10.1093/bib/bbae657

10. Hu, J., Li, X., Coleman, K., Schroeder, A., Ma, N., Irwin, D. J., Lee, E. B., Shinohara, R. T., & Li, M. (2021). SpaGCN: Integrating gene expression, spatial location and histology to identify spatial domains and spatially variable genes by graph convolutional network. Nature Methods, 18(11), 1342–1351. 10.1038/s41592-021-01255-8

11. Kim, J., Rustam, S., Mosquera, J. M., Randell, S. H., Shaykhiev, R., Rendeiro, A. F., & Elemento, O. (2022). Unsupervised discovery of tissue architecture in multiplexed imaging. Nature Methods, 19(12), 1653–1661. 10.1038/s41592-022-01657-2

12. Kinkhabwala, A., Herbel, C., Pankratz, J., Yushchenko, D. A., Rüberg, S., Praveen, P., Reiß, S., Rodriguez, F. C., Schäfer, D., Kollet, J., Dittmer, V., Martinez-Osuna, M., Minnerup, L., Reinhard, C., Dzionek, A., Rockel, T. D., Borbe, S., Büscher, M., Krieg, J., … Bosio, A. (2022).MACSima imaging cyclic staining (MICS) technology reveals combinatorial target pairs for CAR T cell treatment of solid tumors. Scientific Reports, 12(1), 1911. 10.1038/s41598-022-05841-4

13. Kröger, C., Müller, S., Leidner, J., Kröber, T., Warnat-Herresthal, S., Spintge, J. B., Zajac, T., Neubauer, A., Frolov, A., Carraro, C., Freiesleben, S. D., Altenstein, S., Rauchmann, B., Kilimann, I., Coenjaerts, M., Spottke, A., Peters, O., Priller, J., Perneczky, R., … Bonaguro, L. (2024). Unveiling the power of high-dimensional cytometry data with cyCONDOR. Nature Communications, 15(1), 10702. 10.1038/s41467-024-55179-w

14. Lee, E., Chern, K., Nissen, M., Wang, X., Huang, C., Gandhi, A. K., Bouchard-Côté, A., Weng, A. P., & Roth, A. (2023). SpatialSort: a Bayesian model for clustering and cell population annotation of spatial proteomics data. Bioinformatics, 39(Supplement_1), i131–i139. 10.1093/bioinformatics/btad242

15. Levine, J. H., Simonds, E. F., Bendall, S. C., Davis, K. L., Amir, E. D., Tadmor, M. D., Litvin, O., Fienberg, H. G., Jager, A., Zunder, E. R., Finck, R., Gedman, A. L., Radtke, I., Downing, J. R., Pe’er, D., & Nolan, G. P. (2015). Data-Driven Phenotypic Dissection of AML Reveals Progenitor-like Cells that Correlate with Prognosis. Cell, 162(1), 184–197. 10.1016/j.cell.2015.05.047

16. Lin, J.-R., Izar, B., Wang, S., Yapp, C., Mei, S., Shah, P. M., Santagata, S., & Sorger, P. K. (2018a). Highly multiplexed immunofluorescence imaging of human tissues and tumors using t-CyCIF and conventional optical microscopes. ELife, 7. 10.7554/eLife.31657

17. Lin, J.-R., Izar, B., Wang, S., Yapp, C., Mei, S., Shah, P. M., Santagata, S., & Sorger, P. K. (2018b). Highly multiplexed immunofluorescence imaging of human tissues and tumors using t-CyCIF and conventional optical microscopes. ELife, 7. 10.7554/eLife.31657

18. Lin, Y., Wu, T.-Y., Wan, S., Yang, J. Y. H., Wong, W. H., & Wang, Y. X. R. (2022). scJoint integrates atlas-scale single-cell RNA-seq and ATAC-seq data with transfer learning. Nature Biotechnology, 40(5), 703–710. 10.1038/s41587-021-01161-6

19. Long, Y., Ang, K. S., Li, M., Chong, K. L. K., Sethi, R., Zhong, C., Xu, H., Ong, Z., Sachaphibulkij, K., Chen, A., Zeng, L., Fu, H., Wu, M., Lim, L. H. K., Liu, L., & Chen, J. (2023). Spatially informed clustering, integration, and deconvolution of spatial transcriptomics with GraphST. Nature Communications, 14(1), 1155. 10.1038/s41467-023-36796-3

20. Märtens, K., Bortolomeazzi, M., Montorsi, L., Spencer, J., Ciccarelli, F., & Yau, C. (2023). Rarity: discovering rare cell populations from single-cell imaging data. Bioinformatics, 39(12). 10.1093/bioinformatics/btad750

21. Momenzadeh, A., & Meyer, J. G. (2025). Single-cell proteomics using mass spectrometry. Cell Genomics, 100973. 10.1016/j.xgen.2025.100973

22. Singhal, V., Chou, N., Lee, J., Yue, Y., Liu, J., Chock, W. K., Lin, L., Chang, Y.-C., Teo, E. M. L., Aow, J., Lee, H. K., Chen, K. H., & Prabhakar, S. (2024). BANKSY unifies cell typing and tissue domain segmentation for scalable spatial omics data analysis. Nature Genetics, 56(3), 431–441. 10.1038/s41588-024-01664-3

23. Van Gassen, S., Callebaut, B., Van Helden, M. J., Lambrecht, B. N., Demeester, P., Dhaene, T., & Saeys, Y. (2015). FlowSOM: Using self-organizing maps for visualization and interpretation of cytometry data. Cytometry Part A, 87(7), 636–645. 10.1002/cyto.a.22625

24. Vandereyken, K., Sifrim, A., Thienpont, B., & Voet, T. (2023). Methods and applications for single-cell and spatial multi-omics. Nature Reviews Genetics, 24(8), 494–515. 10.1038/s41576-023-00580-2

25. Weigert, M., & Schmidt, U. (2022). Nuclei Instance Segmentation and Classification in Histopathology Images with Stardist. ISBIC 2022 - International Symposium on Biomedical Imaging Challenges, Proceedings. 10.1109/ISBIC56247.2022.9854534

